# Braess’ Paradox in Enzyme Kinetics: Asymmetry from Population Balance without Direct Cooperativity

**DOI:** 10.1101/2025.07.30.667590

**Authors:** Malte Schäffner, Colin A. Smith, Robert Tampé, Helmut Grubmüller

## Abstract

The ATPase ABCE1, a member of the ubiquitous ATP-Binding Cassette protein superfamily, is essential in eukaryotic and archaeal ribosome recycling. It comprises a pair of homologous nucleotide-binding domains each containing a consensus nucleotide-binding site (NBS), where ATP hydrolysis takes place. Each of these sites can be either in an open or closed conformation. Despite this near symmetry, and quite unexpectedly, their hydrolysis kinetics are highly asymmetric. While substitution of the catalytic glutamate (E238Q) in NBSI reduced the overall turnover rate of the ATPase by a factor of two, as one might expect, the corresponding substitution in NBSII (E485Q) shows a so far unexplained tenfold increase. To address this issue, we used Markov models to study how such drastic asymmetry can arise. Specifically, we asked if previously proposed direct allosteric interaction between the two nucleotide-binding sites, such as electrostatic interactions, are required to explain this observation. Indeed, using a Bayesian approach, we found Markov models that quantitatively predict the experimentally observed kinetics as well as additional steady state ATP occupancy data without such direct allosteric interaction. In particular, our results show that the structure-induced property that opening and closing always involves both nucleotide-binding sites suffices to explain the observed remarkable asymmetry. These models can explain the unexpected fast kinetics of the mutant of NBSII in terms of a drastic population shift due to the mutation, which circumvents a kinetic trap state that slows down wild-type kinetics. Our Baysian Markov approach may help to quantitatively explain similar non-intuitive Braess-type kinetics also in other enzymes where chemical/conformation coupling is essential.

## 1 Introduction

The protein ATP-binding cassette subfamily E member 1 (ABCE1)^1,2^ is a member of the ATP-binding cassette (ABC) transporter superfamily. In ABC transporters, ABCE1 homologues^3^ serve as chemo-mechanic energy converters, which drive conformational changes of transmembrane domains, which in turn affect transport of the respective substrate^4^. In this sense, the soluble protein ABCE1, which is void of transmembrane parts, is a prototype of the ‘molecular motor’ of this large and crucial membrane protein family. In fact, based on the very high sequence conservation, being one of the most conserved proteins in evolution, ABCE1 might even be a prototype for a very ancient molecular motor. As an exception within this protein superfamily, ABCE1, in coordination with other translation factors, drives separation of the large and small ribosomal subunit via a drastic displacement of an N-terminal Fe4S4 cluster domain^5–8^, which also involves chemo-mechanical energy conversion.

ACBE1 is mainly composed of two sequentially and structurally highly similar nucleotide binding domains (NBD)^7–14^. Figure 1A shows the structure of ABCE1, highlighting seven highly conserved sequence motifs present in each of the two NBDs, which are required for ATP-binding and hydrolysis, namely the A-loop (Y-loop), Walker-A (P-loop), Q-loop, His switch (H-loop), Walker-B, D-loop, and the ABC-signature motif (C-loop).

**Figure 1.**
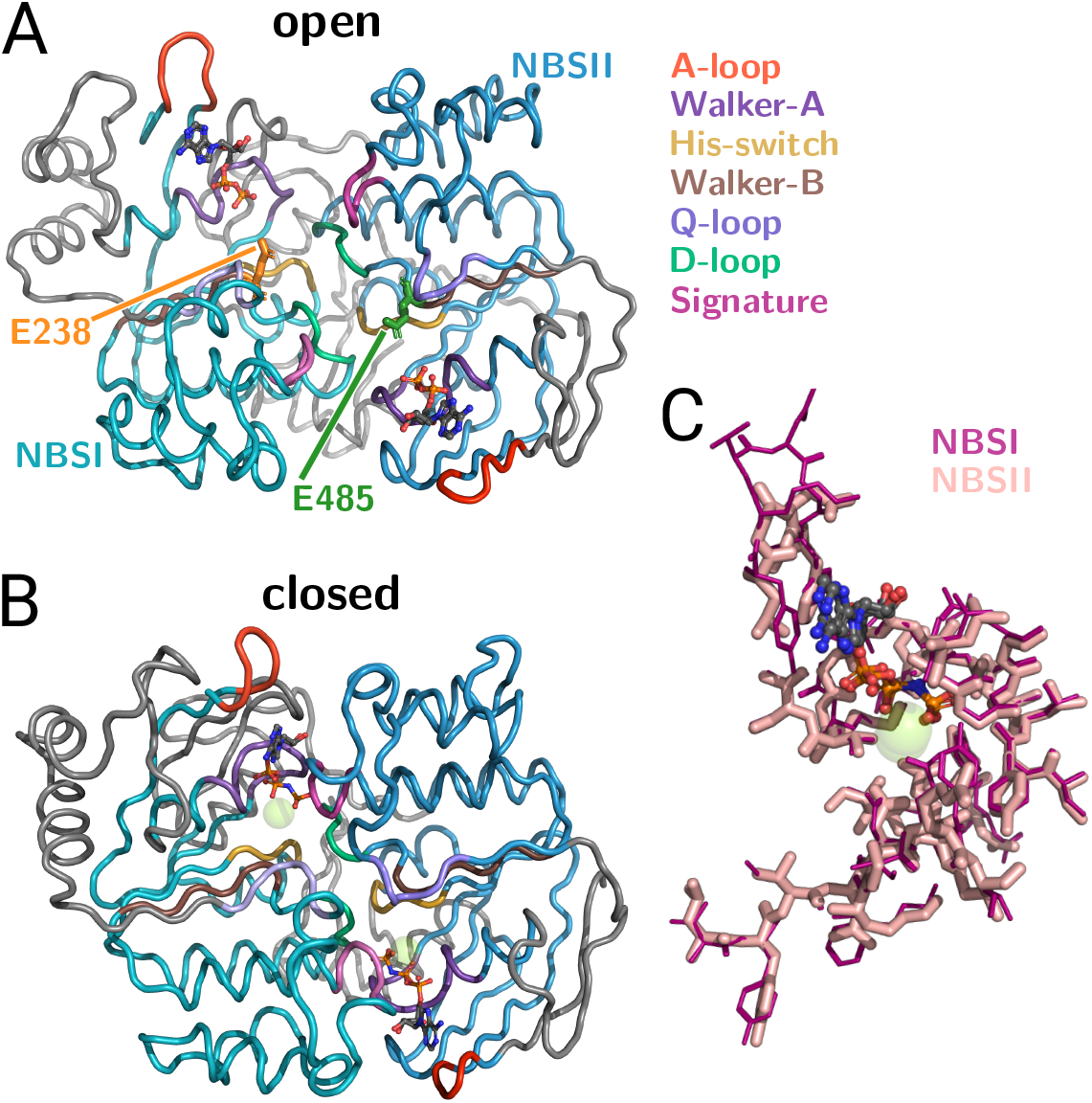
Structure of ABCE1 lacking the FeS domain. (A) Open conformation (pdb code 3BK7^10^) and (B) closed conformation (pdb code 6TMF^10^) in ribbon representation (NBD1: turquoise, NBD2: cyan, all other residues in gray) with bound ligands (ADP, ATP analog AMP-PNP, SF4 and Mg) in ball-and-stick representation. Regions referred to in the main text are indicated by color. (C) Selected regions of ABCE1 in its closed conformation with two bound AMP-PNPs molecules (pdb code 6TMF^8^). Shown is a superposition of the two quasi-symmetric nucleotide-binding sites NBSI and NBSII based on alignment of the seven highly conserved residues of each nucleotide-binding site.

As expected based on the high similarity of the NBDs, ABCE1 also hosts two highly simi lar ATP-hydrolysis competent nucleotide-binding sites, referred to as NBSI and NBSII. Both nucleotide-binding sites are located at the interface between the two anti-parallel oriented NBDs and in fact comprise residues of both NBDs. A structural alignment and superposition of the two nucleotide-binding sites (Figure 1C) further illustrates the high similarity between both nucleotide-binding sites.

Multiple ABCE1 structures have been solved^7–24^, which, taken together, suggest a ligand dependent opening and closing motion of the dimer as the elementary chemo-mechanical process^8^. The specific structure of the dimer further suggests that this opening and closing motion always involves both nucleotide-binding sites, i.e., the two nucleotide-binding sites are dynamically highly coupled such that one cannot be open while the other is closed^25^. This feature will turn out to be crucial for our understanding of the molecular determinants of ABCE1 function.

In addition to the available structural information, bulk and single-molecule experiments on ABCE1 have been carried out to characterize the thermodynamics and kinetics of ABCE1^7,25,26^. In particular, to obtain information on the individual nucleotide-binding sites, two site-selective ATP-hydrolysis defective mutants were generated, and the ATP turnover rates of these mutants were measured and compared to the wild type (WT)^7^. In each of the two mutants, the catalytically active Walker-B motif glutamate of one nucleotide-binding site was changed to glutamine, while that of the other binding was left intact, resulting in drastically reduced ATP catalysis rates of the respective nucleotide-binding site. Mutant E238Q has the modification in NBSI and mutant E485Q in NBSII, allowing the ATP turnover rate of the opposite nucleotide-binding site to be measured separately in each case.

The high similarity between NBSI and NBSII would suggest that each nucleotide-binding site contributes equally to the wild-type ATP turnover rate, such that abolishing one of the nucleotide-binding sites while leaving the other intact should reduce the turnover by a factor of about two compared to the wild type. This reduction has indeed been observed for mutant E238Q; however, and quite unexpectedly, mutant E485Q did not show a similarly reduced ATP turnover. Instead, inactivating NBSII results in a tenfold increased turnover rate^7,26^! Such pronounced asymmetry is in fact unprecedented among all members of the ABC superfamily. For example, MDR1A P-glycoprotein shows a quite symmetric decrease in ATP turnover rate for mutants with identical EQ point mutation in each nucleotide-binding site^27^.

It has been suggested that this striking asymmetry reflects different functions of the two nucleotide-binding sites. Whereas NBSI may control structural changes of the FeS domain resulting in ribosome splitting, NBSII has been suggested to act as a ‘timer’ that triggers dissociation of the 30S ribosomal subunit after ATP hydrolysis, thereby terminating ABCE1 function in ribosome anti-association.^26^

Clearly, such unexpected asymmetric kinetics calls for an explanation. An straightfor ward one is to assume a direct allosteric communication between both nucleotide-binding sites, such that ATP-binding in NBSII directly affects and facilitates ATP-binding or hydrolysis (or both) within NBSI^8,25,26^. This hypothesis would explain the increased ATP turnover rate of mutant E485Q in terms of an increased ATP occupation of hydrolysis-defective NBSII and, thus, by an—on average—higher population of molecules with ‘activated’ NBSI^8^.

In the following we briefly summarize the support for this hypothesis by comparison with other members of the ABC superfamily with homologous NBDs, where some evidence indicates an allosteric pathway that facilitates direct communication between the two nucleotide binding sites.

To investigate a possible allostery in the bacterial exporter Sav1866, a 150 ns molecular dynamics simulation was carried out starting from an AMP-PNP/AMP-PNP crystal structure (notation gives the occupation of NBSI and NBSII, respectively)^28^. To mimic ATP unbinding, the nucleotide-binding site occupations were changed to ATP/apo and the simulations resulted in a hydrolysis-competent state of NBSI.

Based on the simulations, the ATP unbinding from NBSII was suggested to be followed by a series of conformational rearrangements from the Walker-B motif of NBSII through the D-loop of NBDII to the Walker-A motif of NBSI^28^. Although this simulation study suggests the possibility of direct allosteric communication, it has the reverse effect compared to what would be required for ABCE1, in that ATP binding in NBSII results in inactivation rather than activation of NBSI.

A similar communication pathway was proposed based on the comparison of apo/apo and ATP/apo bound crystal structures of bacterial exporter TM287/TM288, where binding of ATP to NBSI affects the stereochemistry of the Walker-A motif of NBSII^29^. However, TM287/TM288 is a heterodimeric ABC exporter and, in contrast to ABCE1, has one nucleotide-binding site—termed degenerate or noncanonical—that lacks critical conserved residues for ATP hydrolysis, thus rendering the nucleotide-binding site ATP-hydrolysis in competent. It is hypothesized that in transporters containing one canonical and one non canonical nucleotide-binding site, the two sites have evolved to perform distinct functions, in contrast to transporters with two canonical sites, where both sites exhibit equivalent functionality^30^. This functional divergence may result in differences in the communication between the nucleotide-binding sites in these distinct transporter types, potentially limiting the applicability of findings from such asymmetric transporters to ABCE1. Additionally, molecular dynamics simulations of TM287/TM288 showed no indication of allostery between the nucleotide-binding sites when started from the ATP/apo structure with ATP added to the second nucleotide-binding site^31^. Detailed structure information under turnover conditions is also available for heterodimeric bacterial exporter TmrAB^32^, albeit without indication of allostery, too.

Finally, in the DNA double-strand break repair protein Rad50 an alanine mutation of the D-loop aspartate resulted in reduced cooperativity of ATP hydrolysis^33^, which indicates an involvement of the D-loop aspartate in the direct allosteric communication between the nucleotide-binding sites. However, neither wild type nor mutants of ABCE1 show any such cooperativity^7^, rendering support along similar lines impossible.

In summary, and given these differences between the homologues, it seems to us that the evidence for—as well as against—direct allosteric communication between the two nucleotide binding sites of ABCE1 is weak. In the absence of further support, here we ask: It is possible to explain all available experimental data of free ABCE1—and particularly the striking turnover rate asymmetry—without recourse to any direct allosteric interactions? And if so, what is the underlying molecular mechanism?

To this end, we describe the chemical reactions (ATP hydrolysis) and conformational transitions of ABCE1 as a Markov model. It has in fact been shown previously that chemical reaction networks are, under certain conditions, equivalent to Markov models^34^. Ac cordingly, we describe the ABCE1 reaction cycle in terms of a set of discrete Markov states (Figure 2A), where each state represents a particular conformational (open or closed) and chemical (empty, ATP-bound, or ADP+P_*i*_-bound) state of ABCE1. Accordingly, transitions between these states represent conformational changes, ligand exchange, or ATP catalysis^35^. In describing these transitions within a Markovian framework, we assume that they are memoryless, i.e., that all transition rates only depend on the current state. Such coarse grained yet thermodynamically consistent description^36,37^ enables us to define the Markov model of ABCE1 such that any direct allostery between the nucleotide-binding sites is excluded *by construction*. If such a model is nevertheless capable of reproducing the asymmetric turnover rates observed for the wild type and both mutants, one would conclude that, indeed, allostery is not required to explain this asymmetric ABCE1 kinetics. Of course, such a result would not rule out direct allosteric interactions between the two nucleotide-binding sites either.

**Figure 2.**
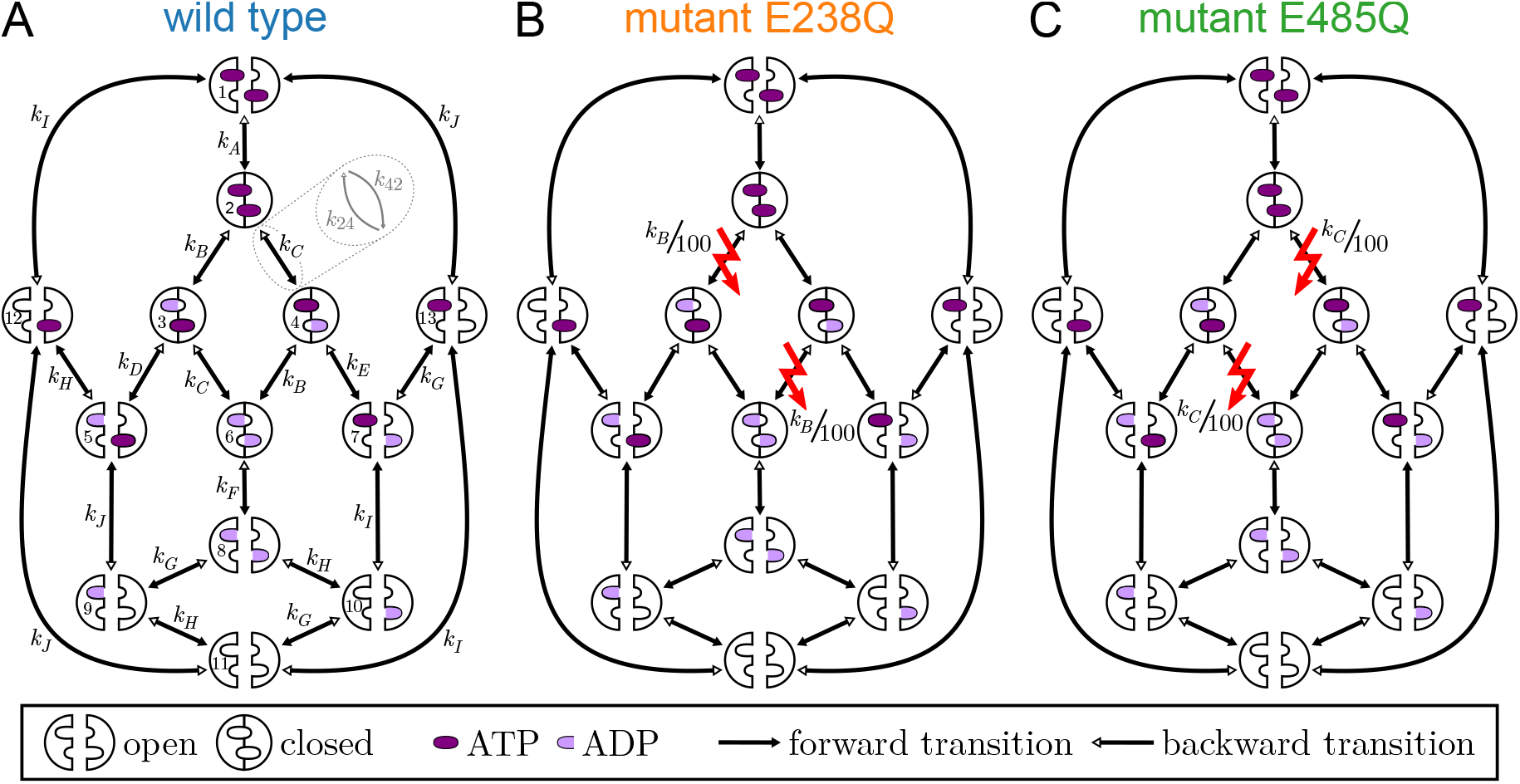
Graph representations of ABCE1 Markov model classes (A) wild type, (B) mutant E238Q, and (C) mutant E485Q. Each ABCE1 symbol represents one Markov state, which are connected by conformational or chemical transitions (arrows) with indicated rate coefficients *k*_*i*_; here rate coefficients that are assumed to be equal are indicated by identical labels (A,B,C…, see also supplementary Table S1). All transitions are reversible; for clarity only the forward rate (solid arrow heads) is annotated. For the two mutants, the reduced ATP hydrolysis rate coefficients of the respective defective nucleotide-binding site are indicated by the red lightning symbols.

To this end, we proceed as follows: First, we only define the ‘topology’ of the Markov model, i.e., its states and its connections. We refer to this model level, which does not include specification of transition rate coefficients between the Markov states, as a Markov model *class*. Each Markov model class, therefore, comprises many specific Markov models, which are defined, in addition, by their transition rate coefficients. Hence we ask if Markov models of this Markov model class can be identified which agree, within experimental uncertainty, with the measured ATP hydrolysis kinetics.

To assess whether such models exist, we employ Bayesian inference to determine the probability of each Markov model given the available measurements^38^. Should many such Markov models with non-negligible probabilities be found, we will group them into sets representing similar molecular mechanisms. In contrast to previous maximum-likelihood based approaches^36,39–41^, this Bayesian approach will also enable us to rank different possible mechanisms according to their probability.

## 2 Results

To test the hypothesis that no direct allosteric communication between the two nucleotide binding sites of ABCE1 is required to explain the asymmetric ATP hydrolysis kinetics of the two mutants, we describe the combined conformational dynamics and ATP binding, hydrolysis and product unbinding as a Markov model, which by construction does not contain such direct interaction. Therefore, if the observed peculiar and counter-intuitive ATP hydrolysis kinetics can be reproduced by the Markov model, we can conclude that these kinetics can be explained without direct allostery.

Note that we aim to explain the asymmetric ATP hydrolysis kinetics as measured by Barthelme et al.^7^ on free ABCE1 in absence of ribosomal subunits and further translational factors. Thus, our models of ABCE1 will also describe free ABCE1 and do not include any ribosome-bound states of ABCE1.

### Markov Model Class of ABCE1

First, we define the states and transitions without specifying particular transition rate coefficients between states. We will refer to this definition as a ‘Markov model class’, and instances of that class with specified transition rate coefficients will be referred to simply as ‘Markov model’. Our goal is to design a Markov model class of ABCE1 that is on one hand minimal in terms of number of states and transitions (to keep the number of parameters low) and on the other hand is complex enough to capture the asymmetry of ABCE1. Furthermore, we aim at a Markov model that is thermodynamically consistent, i.e., that all relevant states are considered, that the concentrations of ATP, ADP and P_*i*_ are properly taken into account, and that detailed balance is obeyed. Detailed balance requires that all transitions are bidirectional.

Figure 2A shows the Markov model class of the wild type. Here we used the following assumptions to determine its number of states, their association with conformational and occupational states of ABCE1, and the transitions between them.

First, we assumed that two conformational states—an open and a closed state—of the 9 nucleotide-binding sites suffice to describe ATP hydrolysis of free ABCE1. These two states are well known from X-ray structures of free ABCE1 occupied by ADP showing ABCE1 with nucleotide-binding sites in an open conformation^7,9,10^ and from cryo-EM structures of ABCE1 bound to the small ribosomal subunit showing ABCE1 with nucleotide-binding sites in a closed conformation^8,14,42^.

In addition to the open and the closed state, cryo-EM structures of pre-splitting 70S ABCE1 and 80S-ABCE1 complexes show the nucleotide-binding sites of ABCE1 in an intermediate or half-open conformation^11,12,16–20^. Additional support for such an intermediate state of nucleotide-binding sites comes from FRET efficiency distributions of dyes attached to the two NBD-parts of each nucleotide-binding site, which suggest at least three distinct states^25^. On a functional level, the intermediate state of ABCE1 is considered necessary for regulation of ribosome separation^8^. In the absence of evidence for a role of this intermediate state in the ATP hydrolysis of free ABCE1, we tentatively assumed that an intermediate third conformational state is not required to explain the kinetic asymmetry of the mutants.

Second, we assumed that both nucleotide-binding sites adopt the same conformation at all times—either both are closed or both are open. This assumption is supported by the fact that all free, 80S- and 30S-bound ABCE1 structures known to us show both nucleotide binding sites in identical conformations^7–12,14–21,24^. We note that cryo-EM structures of 43S and 48S late initiation complexes show NBSI in the intermediate conformation and NBSII in the closed conformation bound with GMP-PNP/GMP-PNP^21,24^ or ADP/ATP^22,23^, respectively. Also, differences in FRET efficiencies corresponding to the two nucleotide binding sites indicate a certain degree of conformational independence of the nucleotide binding sites^25^, but no direct measurement of the correlation between the conformational states of the two nucleotide-binding sites exists to date. Whereas such evidence may suggest a certain degree of conformational independence between the nucleotide-binding sites, we decided to first test if the asymmetric ATP hydrolysis kinetics can be reproduced without assuming such independence.

Third, we assumed that ABCE1 closes only if both nucleotide-binding sites are occupied by a nucleotide. Indeed, all known closed structures^8,14,42^ and intermediate/closed structures^13,21–24^ of sufficient resolution do show nucleotides in both nucleotide-binding sites. Additional experimental evidence comes from mutants of ABCE1 that are unable to bind nucleotides in one or both nucleotide-binding sites^26^. These mutants are also unable to split ribosomes, which requires closing of both nucleotide-binding sites^8,14,42^, and, consequently, binding of two nucleotides is necessary for ABCE1 to adopt a closed state and split ribosomes^26^. Further, in absence of nucleotides, FRET-experiments showed that NBSII remains in an open conformation^25^. This assumption lead us to omit closed conformations with empty nucleotide-binding sites from our Markov model class and, furthermore, reduces the number of transitions between Markov states representing open and closed conformations of ABCE1.

The next two assumptions define the transitions between Markov states of equal conformation.

The fourth assumption is that ligand exchange occurs only in the open conformation, which is sterically plausible because X-ray structures^7,9,10,15^ of the open conformation show that the distance between both NBDs, termed nucleotide-binding cleft, is large (10 to 14 Å^10^). In contrast, in all structures of ABCE1 in the closed state, two AMP-PNP are trapped between the two NBDs keeping them in close proximity without notable nucleotide binding cleft^8,14^.

The fifth assumption is that ATP hydrolysis occurs only in the closed conformation, which, too, is sterically plausible from structures because in the closed conformation all residues required for ATP hydrolysis are in contact with bound AMP-PNP^8,14^. In the closed state, the signature motif and D-loop of the opposite NBD complement the binding site and coordinate the *γ*-phosphate. In the open conformation, the large nucleotide-binding cleft positions these two motifs hydrolysis too far from a bound ATP to establish interactions required for ATP hydrolysis^7,9,10,29,43^. As a result ATP hydrolysis is inhibited in the open conformation.

The sixth assumption is that there is no allosteric communication between the nucleotide binding sites. This is the central assumption to test our hypothesis that this sort of allostery is not necessary for the asymmetric ATP hydrolysis kinetics of ABCE 1mutants. On a technical level this means that the transition rate of one type of transition, e.g., ATP binding to NBSII, are independent from the occupation of the opposite nucleotide-binding site and, thus, these rates occur multiple times in the model as indicated by identical labels *k*_*A*…*J*_ in Figure 2A.

The last assumption, more technical in nature, aims at reducing the complexity of the Markov model. To this aim we assume that ADP- and P_*i*_ exchange can be described by only one effective transition, which implies that the respective on- and off-rates describe the long-timescale kinetics of the two subsequent binding and unbinding events. Thus, our model does not distinguish between ADP-, P_*i*_-, and ADP+P_*i*_-bound states and considers for each conformational state and each nucleotide-binding site of ABCE1 an empty, an ADP+P_*i*_ bound, and an ATP-bound binding state. Accordingly, we will henceforth refer to ADP and P_*i*_ exchange collectively as ADP exchange. This assumption is consistent with the most recent model of ABCE1-driven ribosome separation, which also combines ADP and P_*i*_ release into one transition^8^. Furthermore, this assumption will turn out to be computationally beneficial, as it reduces the number of distinct combinations of occupations from 5^2^(apo, ATP, ADP+P_*i*_, ADP, P_*i*_) to 3^2^(apo, ATP, ADP), thereby reducing the number of Markov states—and, hence, the dimension of the search space—by over twofold.

Recent structural studies of bacterial exporters TmrAB^32^ and TAP1/2^44^ have shown that in these transporters P_*i*_ dissociates from the protein prior to the opening of the nucleotide binding sites and the subsequent release of ADP. If this mechanism is also applicable to ABCE1, our results would remain valid, albeit with a revised interpretation of the transitions. In this case, the effective transition for ADP+P_*i*_ binding is unchanged, while the ADP+P_*i*_ unbinding transition would be redefined as only ADP unbinding. The current opening transition would then also encompass P_*i*_ release.

Taken together, the above assumptions result in the Markov model class of ABCE1 shown in Figure 2A with 13 states and 10 forward and 10 backward transition rate coefficients as free parameters. Supplementary Table S1 lists all transition rate coefficients along with the corresponding conformational changes, chemical reactions, or ligand exchanges. The Markov model classes of mutants E238Q and E485Q (Figure 2B and C) are derived from the wild-type class by reducing the transition rate coefficients of ATP hydrolysis (*k*_*B*_ and *k*_*C*_) and synthesis (*k*_−*B*_ and *k*_−*C*_) of the mutated nucleotide-binding site by a factor of 100. This reduction describes the fact that the mutation only renders the nucleotide-binding site ATP-hydrolysis defective rather than completely blocking it^7,26,27^. Further, this reduction implements as an eighth assumption that the point mutations affect only the ATP catalysis of the respective nucleotide-binding site, leaving catalytic rates and binding affinities of the other nucleotide-binding site unaffected.

Finally, we incorporated detailed balance into the Markov models, which reduces the number of parameters by three^45^. Rather than implementing proper restrictions on the transition rate coefficients, however, we derived the latter from assigning a free energy to each Markov state, which we consider more intuitive and straightforward. As a result, our Markov models are parametrized by 17 parameters, four free binding energy differences (ATP and ADP binding for each nucleotide-binding site), one free energy difference associated with the conformational change, two free energy differences between ATP and ADP bound in the closed state (one for each nucleotide-binding site), and one free energy barrier for each transition A,B,….,J (see Figure 2A). To convert between free energies and transition rate coefficients, we used transition state theory^46^, which states that

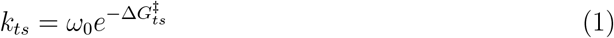

where *k*_*ts*_ is the transition rate coefficient from state *s* to state *t*, 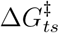 is the free energy barrier between state *s* and state *t*, and *ω*_0_ is the attempt frequency. Here, the particular choice of *ω*_0_ is irrelevant, as any change of *ω*_0_ can be absorbed into the barrier heights; to allow for intuitive interpretation of the barrier height, we have chosen 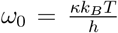 with *κ* the transmission coefficient (hereafter set to one), *k*_*B*_ the Boltzmann constant, *T* the temperature, and *h* the Planck constant.

These Markov models will subsequently serve to calculate and compare the measured kinetic observables, i.e., limiting ATP turnover rate and Michaelis-Menten constant.

Markov Models Agree With Asymmetric ATP Hydrolysis Kinetics To determine whether Markov models exist that agree with the measured ATP hydrolysis kinetics^7^, we used Bayesian inference. Specifically, for each Markov model *M*, its posterior probability is calculated via

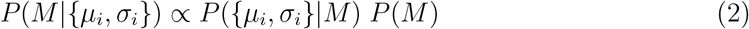

for the six measured values *µ*_*i*_ (3 ATP turnover rates and 3 Michaelis-Menten constants for wild type, mutant E238Q, and mutant E485Q) and their respective experimental uncertain ties *σ*_*i*_. Assuming a Gaussian error model for the experiments, the likelihood^38^ reads

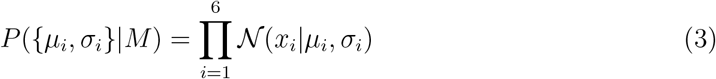

where *N* is a normalized Gaussian and *x*_*i*_is the kinetic observable as calculated from the specific Markov model under consideration. As prior *P*(*M*) a product of uniform priors of the free energies was chosen with physics-motivated upper and lower limits, depending on the type of the particular transition (see Methods). This choice of the prior implies a log-uniform prior for the respective transition rate coefficients and transition times. For posterior sampling a Markov-chain Monte Carlo algorithm^47^ was used with multiple chains to assess convergence of Bayes-sampling.

To check whether the obtained distribution of Markov Models results in the expected distribution of experimental observables centered at the respective measured values, all six kinetic observables were calculated for each Markov Model (see Methods) and shown as histograms in Figure 3. As can be seen, both limiting ATP turnover rate *k*_*cat*_ (Figure 3A) and Michaelis-Menten constant *K*_*M*_ (Figure 3B) agree well with the measured values and are distributed following the experimental uncertainties *σ*_*i*_, indicating correct and sufficiently converged Bayes-sampling of the Markov models.

**Figure 3.**
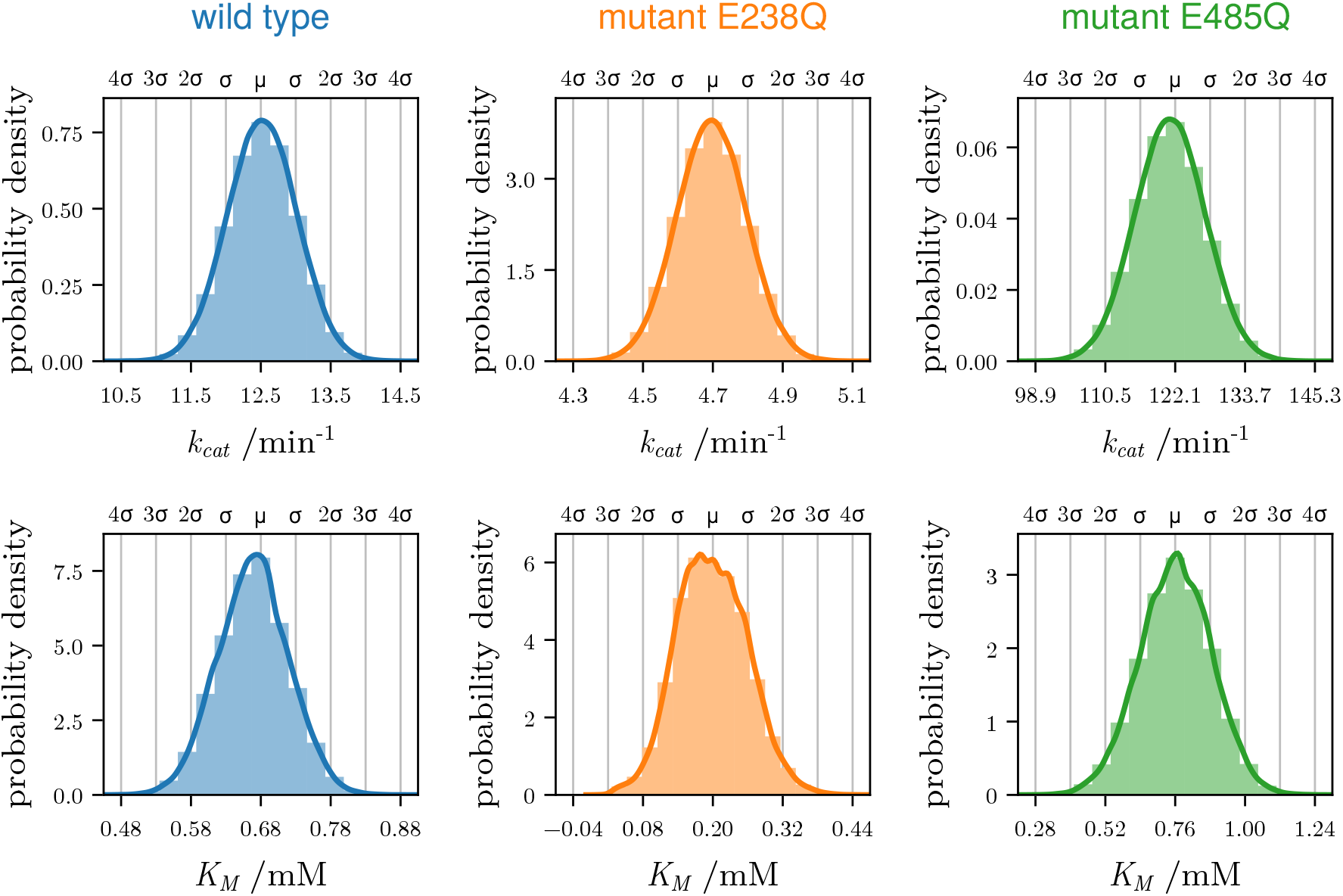
Posterior probability distributions of Michaelis Menten constants. Shown are histograms and fitted kernel densities (solid lines) of *k*_*cat*_ (top row) and *K*_*M*_ (bottom row) for wild type (left), E238Q (mid), and E485Q (right), as obtained from Bayes-sampling of the respective Markov models. The annotations on the top of each plot show the measured values^7^ *µ* and experimental uncertainties (in units of standard deviation *σ*).

A sufficiently converged Bayes-sampling is further supported by major overlap between multiple Markov-chain Monte Carlo runs for all free energy parameters with minor deviations of individual runs, for example, for barrier 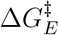 of the conformational change with ATP/ADP occupations for values ⪆ 38 kT (see supplementary Figure S1).

Next, we asked whether there are any Markov models which agree with all six measured kinetic observables and if so, how many. We found that 9.3 % of all sampled Markov models agree with all six measured observables within experimental uncertainty *σ*_*i*_ and, hence, reproduce the observed striking asymmetry.

We conclude that, at least within our Markovian framework, no direct communication between the two nucleotide-binding sites is required.

### Lopsided Wild-type Populations Enable Asymmetric ATP Hydrolysis Kinetics

Next, we investigated the mechanism by which the asymmetric ATP hydrolysis kinetics arises in our Markov models. To this end, we identified in each Markov model those reaction cycles which contribute most to the ATP hydrolysis. Here, reaction cycles are defined as closed sequences of Markov states which involve one hydrolysis step, and the contribution *θ* to ATP hydrolysis was quantified by the probability net flux Θ, i.e., the difference of population multiplied by the transition rate coefficient between two states, through the respective cycle relative to the total ATP hydrolysis rate of that Markov model. For instance, the sequence of states ‘1-2-3-5-12’ corresponds to the hydrolysis of ATP in NBSI with ATP bound in NSBII. Hence, the sum of the contributions *θ* of all hydrolyzing reaction cycles is the total hydrolysis rate.

We used these contributions to classify all sampled Markov model that agreed with all measurements within experimental uncertainty into ‘reaction types’. To this end, we considered only ‘dominant’ reaction cycles with *ϑ* ≥ 20 % at saturating concentrations of ATP, and grouped them accordingly. Importantly, the Bayesian sampling according to the posterior (Equation 2) enabled us to calculate the probability of each reaction type as the normalized count of Markov models of that type.

Further support for this grouping comes from an analysis of the distributions of the net fluxes separated by ATP hydrolysis transitions. As shown in supplementary Figure S2A, most of these distributions are multimodal, and our grouping into reaction types decomposes them into monomodal distributions (supplementary Figure S2B-D).

Strikingly, a total of almost 500 different reaction types were determined, highlighting the complexity of a complete reaction kinetics analysis of even such a comparingly simple biomolecule. Figure 4 shows one randomly selected example for each of the three most probable reaction types (‘A1’, ‘A2’, ‘B’), which, taken together, account for 40.4 % of all sampled Markov models and, hence, deserve closer analysis. In the Figure, the net flux Θ is indicated via the width and color of each arrow (transition) and the size and color of each state (vertex) indicates the steady state probability, i.e., the fraction of enzymes expected to be in each state, henceforth referred to as the population. Dominant reaction cycles are drawn solid, all others are dashed.

**Figure 4.**
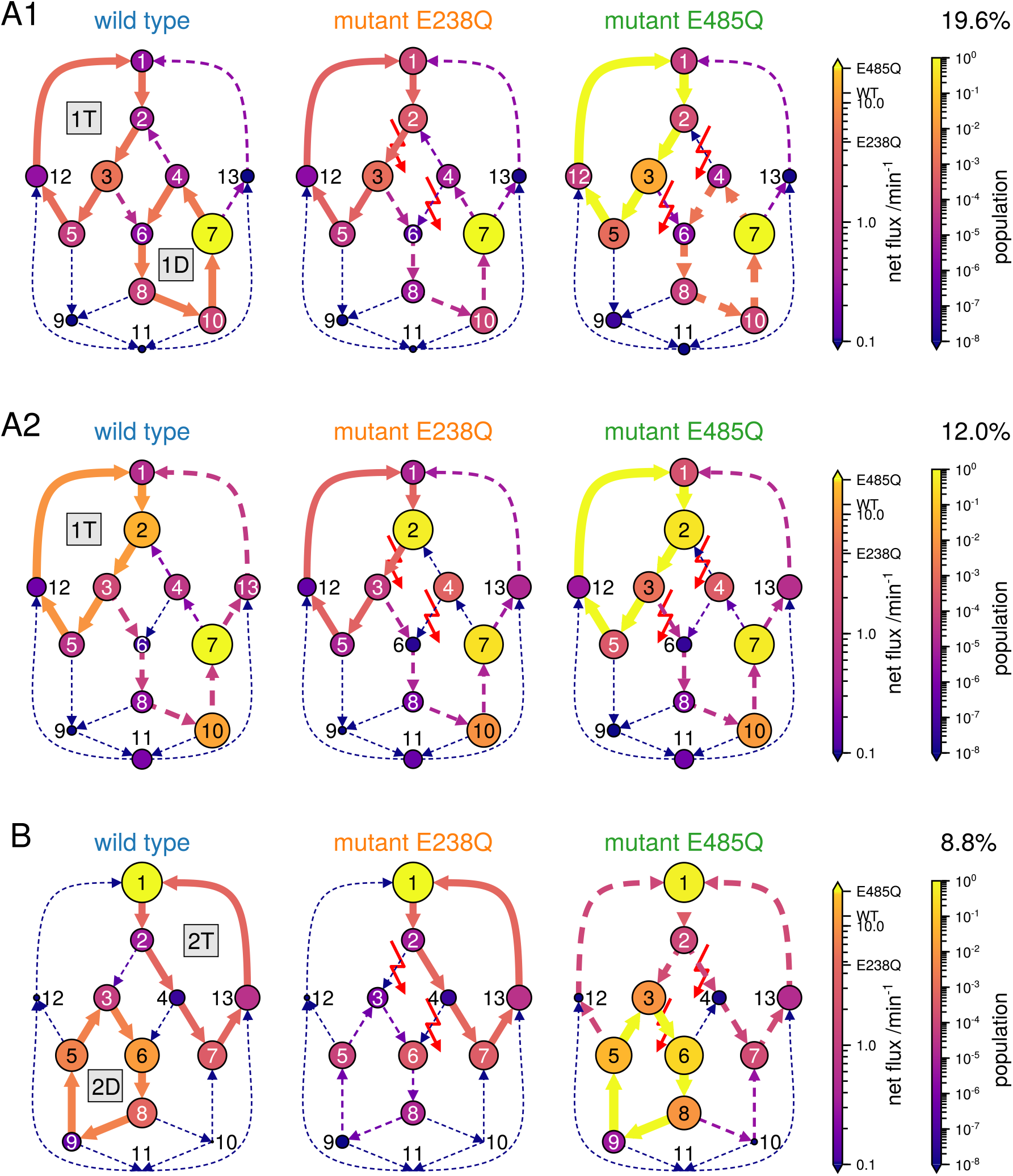
The three most probable reaction types. Their posterior probabilities indicated as percentages; Markov states (circles) are defined as in Figure 2 and their population is indicated by size and color; similarly, the net fluxes through the transitions (arrows) are indicated by line widths and colors. Dominant reaction cycles (defined in the text) are labeled (‘1T’, ‘1D’,…) and are indicated by solid lines, otherwise dashed. For the two mutants, the reduced ATP hydrolysis rate coefficients of the respective defective nucleotide binding site are indicated by the red lightning symbols.

We use the most probable reaction type A1 (Figure 4, top row) to explain a plausible mechanism underlying the asymmetric ATP hydrolysis kinetics (cf. also Figure 2). In this reaction type, mainly two cycles contribute to the overall ATP turnover rate in the wild type. In the upper left cycle (denoted by ‘1T’), hydrolysis occurs in NBSI, while NBSII is ATP-loaded. In the lower right cycle (‘1D’), hydrolysis also occurs in NBSI, with NBSII being ADP-loaded. Thus, unexpectedly, in the dominant reaction type A1, by far most of the ATP hydrolysis is performed by NBSI. Although the net fluxes of cycles 1T and 1D are similar, their kinetics is very different. Compared to cycle 1T, the population 1D states is much (100-1000x) higher which is compensated by lower rates. In fact, almost all of the total population of the system is concentrated in state 7, i.e., ABCE1 is open most of the time, with NBSI loaded with ATP and NBSII with ADP. As will become important below, this population asymmetry between the two cycles is largely controlled by the ATP catalysis rates of NBSII, which creates a net flux from state 3 to 6 and thus acts as a ‘drainage’ of cycle 1T into cycle 1D. As the E485Q mutation hinders ATP catalysis in NSBII, it drastically reduces this drainage. Overall, the observed ATP hydrolysis rate does not simply result from one reaction cycle, but arises from a combination of several cycles and a fine-tuned population balance between these.

This balance is markedly shifted by the E238Q mutation, which eliminates ATP hydrolysis in NBSI almost entirely (Figure 4, top middle). Despite this 100-fold reduction of NBSI, cycle 1T is still active and NBSI is still the main contributor to the overall ATP turnover rate. This counterintuitive response is explained by an increased population of the states upstream of ATP hydrolysis (states 1 and 2), where both NBSs are ATP-loaded. Essentially, this population ‘piles up’ due to the subsequent larger barriers. The strength of this particular effect — as well as precisely which upstream states are being affected — depends on the particular choice of the individual transition rate coefficients. In contrast, cycle 1D is largely suppressed. Here the population of upstream states drains from state 4 into state 2, further contributing to the population increase of upstream states of cycle 1T, thus contributing to the only slight decrease of cycle 1T and the two-fold ATP turnover rate.

The mutation E485Q, in contrast, does not directly affect transitions within the 1T and 1D cycles, but the transitions responsible for the drainage between cycles. As a result, the drainage of cycle 1T via hydrolysis in NBSII is largely suppressed (Figure 4, upper right), resulting in a ca. 10 to 20-fold higher population of this cycle. It is this drastic population shift that causes the nearly 10 to 20-fold increase of the ATP turnover within cycle 1T, despite its transition rate coefficients being unchanged relative to the wild type.

The second most probable reaction type, A2 (Figure 4, middle row), is very similar to reaction type A1. The main difference is that cycle 1T is the sole dominant contributor to the overall ATP turnover rate in the wild type. Still, as with reaction type A1, cycle 1T is not highly populated in the wild type, enabling a large population shift in mutant E485Q, when the drainage from cycle 1T is largely suppressed.

For the second most probable reaction type (Figure 4, bottom row), the mechanism is similar to reaction type A1, but with cycle ‘2D’ taking over the role of 1T and cycle ‘2T’ that of 1D. As a result, the effects of the mutations are also switched. In particular, in reaction type B mutant E238Q now impacts the drainage between the cycles, whereas mutant E485Q impacts cycles 2D and 2T directly. However, the switched roles still lead to the same effect of the mutations on overall ATP turnover. For mutant E238Q, and in contrast to mutant E485Q in reaction type A1 and A2, no large population shift is observed because most population is already located in source cycle 2T of the drainage. Instead, this drainage (from cycle 2T to cycle 2D) is interrupted resulting in suppression of cycle 2D thereby halving the overall ATP turnover rate. For mutant E485Q, an increase in the upstream states (3 and 5) of ATP hydrolysis in NBSII is observed, similar to mutant E238Q in reaction type A1 and A2. However, the effect is even stronger due to the particular choice of the individual transition rate coefficients.

Next, we asked what causes the asymmetric population distribution between the cycles.

To answer this question, Figure 5 shows, for each Markov state separately, the population distribution of all sampled Markov models ignoring their particular reaction type. Clearly, for the wild type, for over 90.6 % of all sampled Markov models, one of the states 1, 6, or 7 is by far the most populated one (≥ 90 % population), leaving only very low populations for all other states.

**Figure 5.**
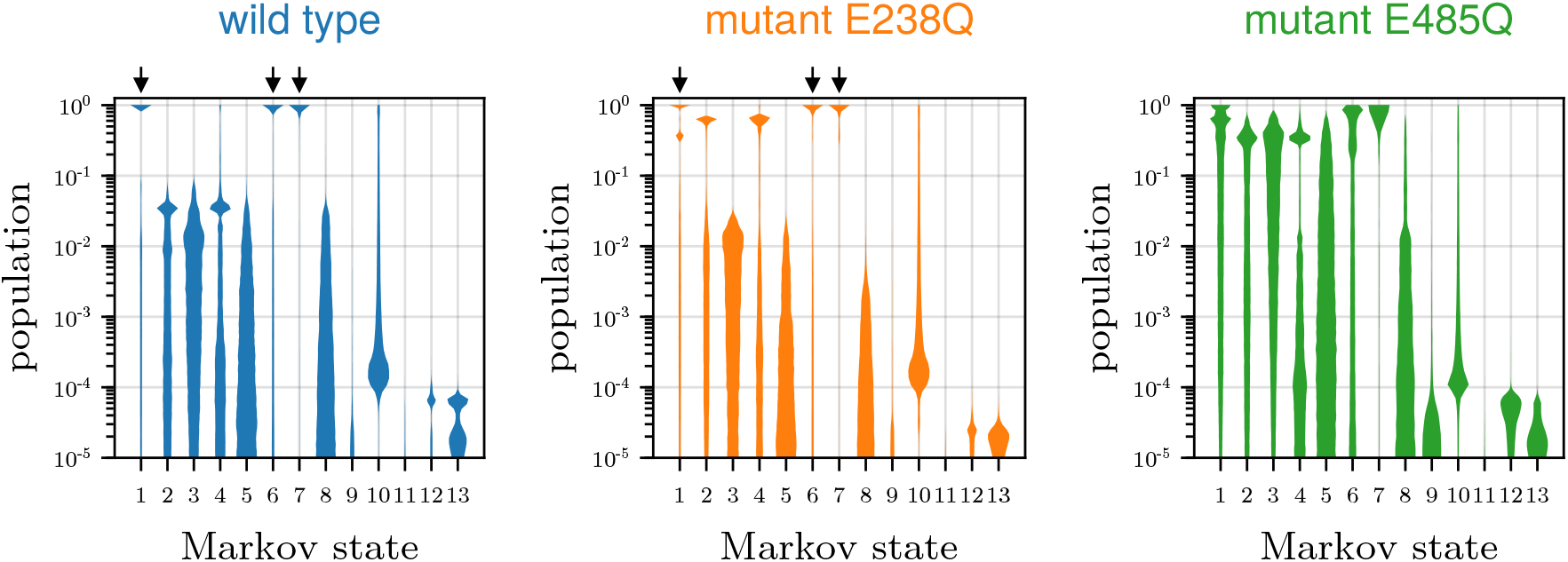
Posterior probability distributions of populations for each Markov state. Shown are distributions of the steady-state populations at conditions of limiting ATP-turnover rate ([ATP] = 100 mol L^−1^) for wild type (left), mutant E238Q (mid), and mutant E485Q (right). Distributions are based on Markov Model samples of posterior 2. Arrows highlight Markov states in which population is concentrated.

This finding is remarkable, because it is not obvious why concentrating nearly all populations in one Markov state should be required to achieve the observed ATP turnover asymmetry between the two mutants.

We speculate that the prevalence of Markov models with population dominance may be due to an ‘entropic factor’, i.e., that for a single state dominated Markov model a larger region of parameter space yields agreement with the experimental data than for Markov models with several markedly populated states. Accordingly, since that latter may require more stringent fine tuning of their parameters, single state dominated Markov models would be ‘easier’ to find during sampling, and may also be more robust. Such robustness of single state dominated Markov models has interesting evolutionary implications.

What property singles out states 1, 6, and 7 from all other states? Are these Markov states perhaps rate limiting for the overall turnover rate, such that the flux through the network is largely dictated by the population of these states? To test this idea, Figure 6A shows the conditional probability for each Markov state to be the most populated one given the rate-limiting Markov state. Here, following the usual convention^48^ the rate-limiting state was defined as the state for which a change of free energy has the largest effect on the overall ATP turnover rate (i.e., the highest degree of ‘thermodynamic’ rate control). Indeed, in all cases the most populated state is also the rate-limiting state of the Markov model.

**Figure 6.**
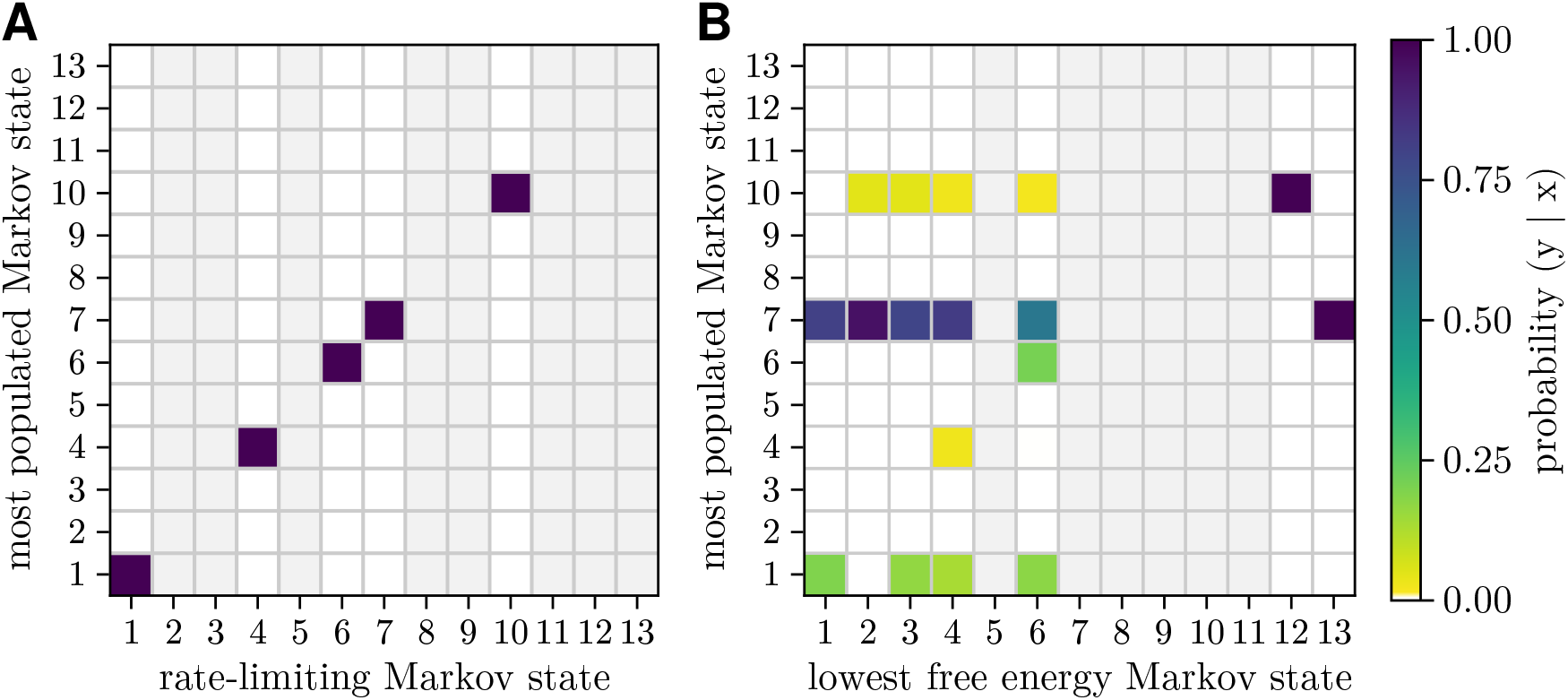
Comparison of the characteristics of wild-type Markov states. Shown are the conditional probabilities (color) of each Markov state to be the most populated, given either (A) the rate-limiting Markov state or (B) the Markov state with the lowest free energy. The column is left empty (gray) for states that were not a rate-limiting or lowest free energy Markov state in any of the Markov models.

To assess whether this agreement can be attributed to the thermodynamic properties of these states, i.e., whether the most populated state corresponds to the lowest free energy state, Figure 6B shows, in a similar manner as Figure 6A, the conditional probability for each Markov state to be the most populated one, now given the lowest free energy state. As can be seen, and in contrast to the rate-limiting state, almost no correlation is seen between the most populated Markov state and the lowest free energy state. This finding strongly supports the notion that the most populated states are highly enriched due to ‘kinetic traps’ and thereby enable relatively high turnover rates despite slow transition rates.

To obtain further insights into why a particular state is the rate-limiting or most populated state, also rate-limiting transition states^49^ and transition rate coefficients (commonly termed sensitivity^50,51^) were computed (shown in supplementary Figure S3). However, no marked correlations were found between most populated state and the rate-limiting transition state. Furthermore, while a transition rate coefficient of an outgoing transition of the most populated state is the rate-limiting transition rate coefficient in 94.8 % of our Markov models, the remaining 5.2 % show that this is neither a necessary nor sufficient condition for determining the most populated state. Both analyses underscore that these ‘kinetic traps’ are due to a more complicated balance between several transition rate coefficients.

On the more technical side, two cautionary notes are in order. First, because the exact definition of ‘dominant reaction cycles’ depends on the particular choice of a threshold, so does the above classification into reaction types. Figure 7 quantifies this dependence and the resulting pattern of dominant cycles (insets). As can be seen, the three reaction types singled out (left insets) and discussed above dominate for a rather wide range of threshold values between 0.05 and 0.25, hence providing justification for their choice. For larger threshold values, Markov models with multiple reaction cycles of similar net flux are characterized by the absence of any dominant reaction cycles. This insensitivity means that models with different reaction cycles are classified into the same reaction type (two top right insets), such that they are not properly distinguished.

**Figure 7.**
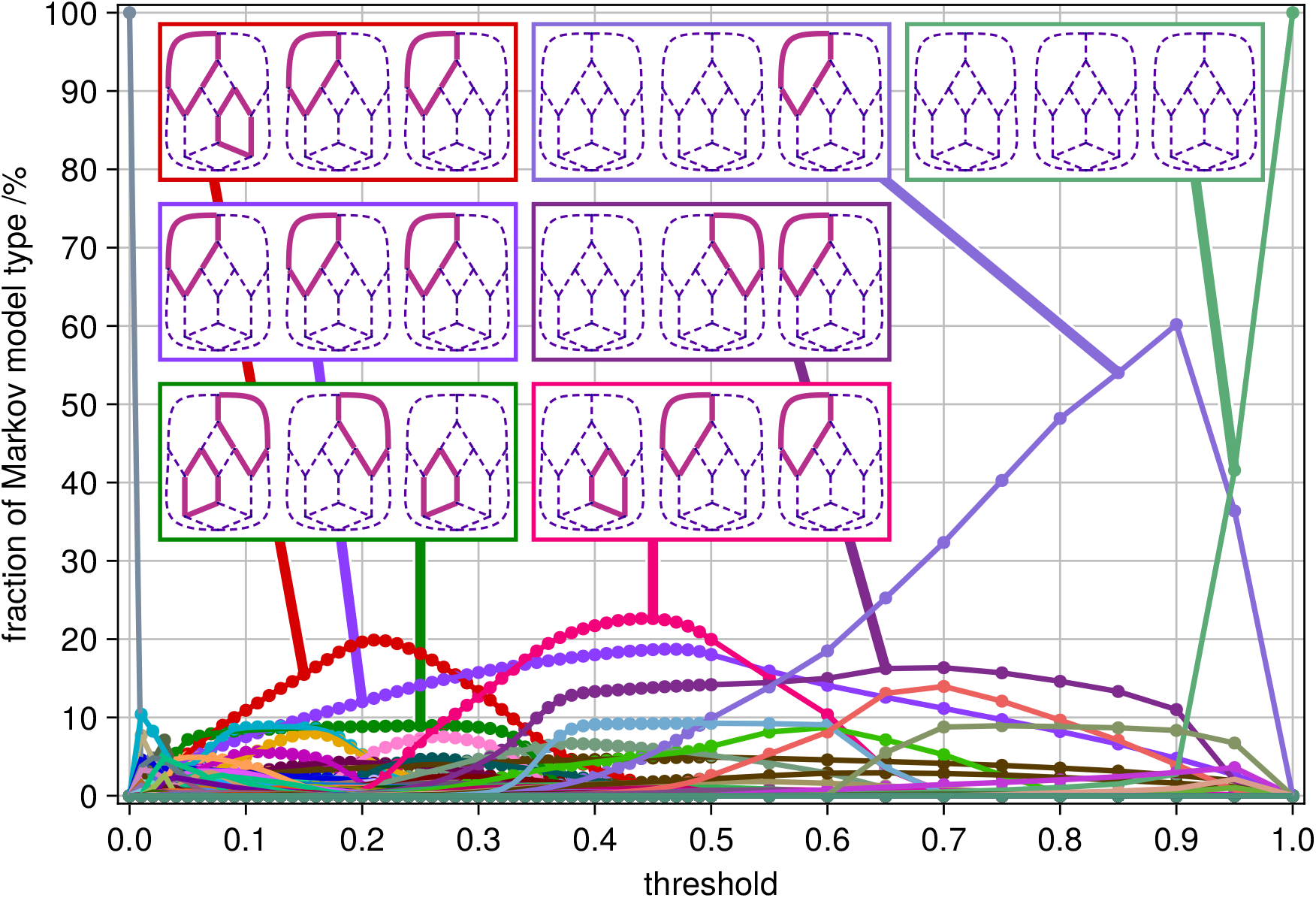
Dominant reaction types vary with chosen threshold. Shown are the posterior probabilities of the most prevalent reaction types (colored lines) for threshold values between 0 and 1. The insets indicate the respective dominant cycles as in Figure 4

Second, it is unknown how much the ATP catalysis rates are reduced by the mutant. Our choice of a factor of 100 is motivated by ATP turnover measurements^7,26,27^, which, most likely, provide only an upper bound. To assess the robustness of our findings against changes of this reduction, we changed it by one order of magnitude in both directions and repeated above Bayesian posterior sampling and reaction type identification protocol. Similar results were obtained; in particular, for a 1000-fold reduction, the same most probable reaction types (B and A1) are seen. For a 10-fold reduction, the kinetic asymmetry was qualitatively reproduced, although, less pronounced in all sampled Markov models, and reaction type A1 was the most probable reaction type (99.9 %). In summary, these results demonstrate that the reaction type we identified as most probable are robust against the experimental uncertainty of how much the E238Q and E485Q mutations actually reduce the ATP catalysis rates.

### Cross-Validation With Ligand Occupancies

Finally, to test for possible overfitting by our Bayesian approach, we checked how well the singled out Markov models agree with new experimental data that have not been used for calculation of the posterior and, hence, for the Bayes-sampling. To this end we used measured ATP and ADP occupancies, for which two different experimental datasets are available—one taken under steady-state conditions^7^ and one measured under non-equilibrium/single-turnover conditions^26^.

Figure 8 shows the steady-state ATP occupancies of Barthelme et al.^7^ as red lines together with posterior ATP and ADP occupation probability densities derived from the Bayesian Markov sample, assuming a steady state. As can be seen, there are Markov models (about 0.07 % of all sampled models) that agree with these new measurements within experimental uncertainty. Although the range of occupancies predicted by our Markov models is too broad to actually predict these experiments, they are consistent with this independent data.

**Figure 8.**
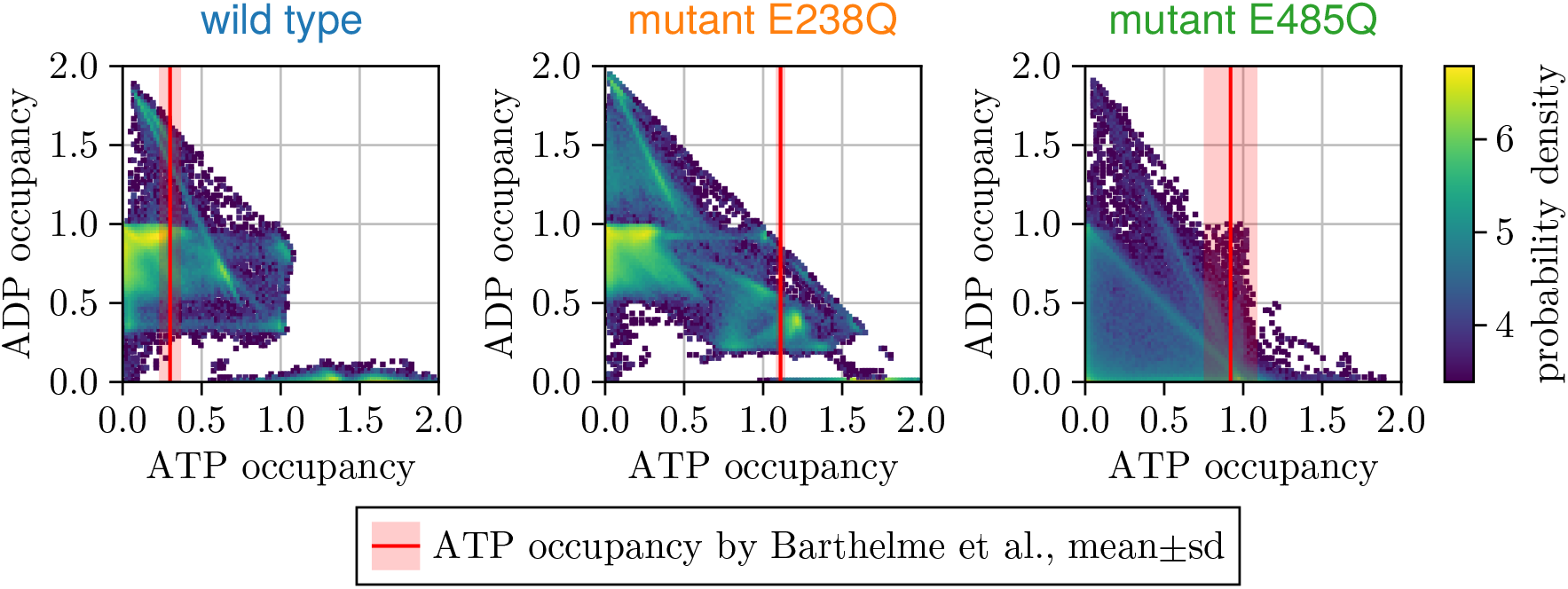
Ligand occupancies. Joint posterior probability distributions of steady-state ATP and ADP occupancies are shown by color for wild type (left), mutant E238Q (middle) and mutant E485Q (right); the measured ATP occupancies^7^(red line) and experimental uncer tainty (shaded, red) are shown for comparison.

The second occupancy measurement was performed by Nürenberg-Goloub et al.^26^ under non-equilibrium conditions. Specifically, a very large initial ABCE1 to ATP ratio (1:2) was used, which results in quickly decaying substrate concentrations even during the short 30 s reaction time of the experiment, so that our steady-state assumption is no longer valid. We therefore resorted to numerical integration of the master equation 4 (see Methods) to model this experiment. Figure 9A compares mean and standard deviation of the measured occupancy distributions by Nürenberg-Goloub et al.^26^ (pink crosses) with the occupancies calculated from the Markov models. Disappointingly, whereas our Markov model sample is consistent with the measured ATP concentrations, much fewer then in the experiments are hydrolyzed during the initial 30 s, pointing to too low hydrolysis rates in essentially all of our Markov models. Also, those Markov models that do hydrolyze ATP within 30 s spend too little time in ADP-bound states to reach the measured occupancies. The fact that this second experiment was performed on different mutants (E → A instead of E → Q) might explain part of this discrepancy; however, because the effect of this mismatch on both NBSs should be rather similar, we do not consider it a satisfactory explanation.

**Figure 9.**
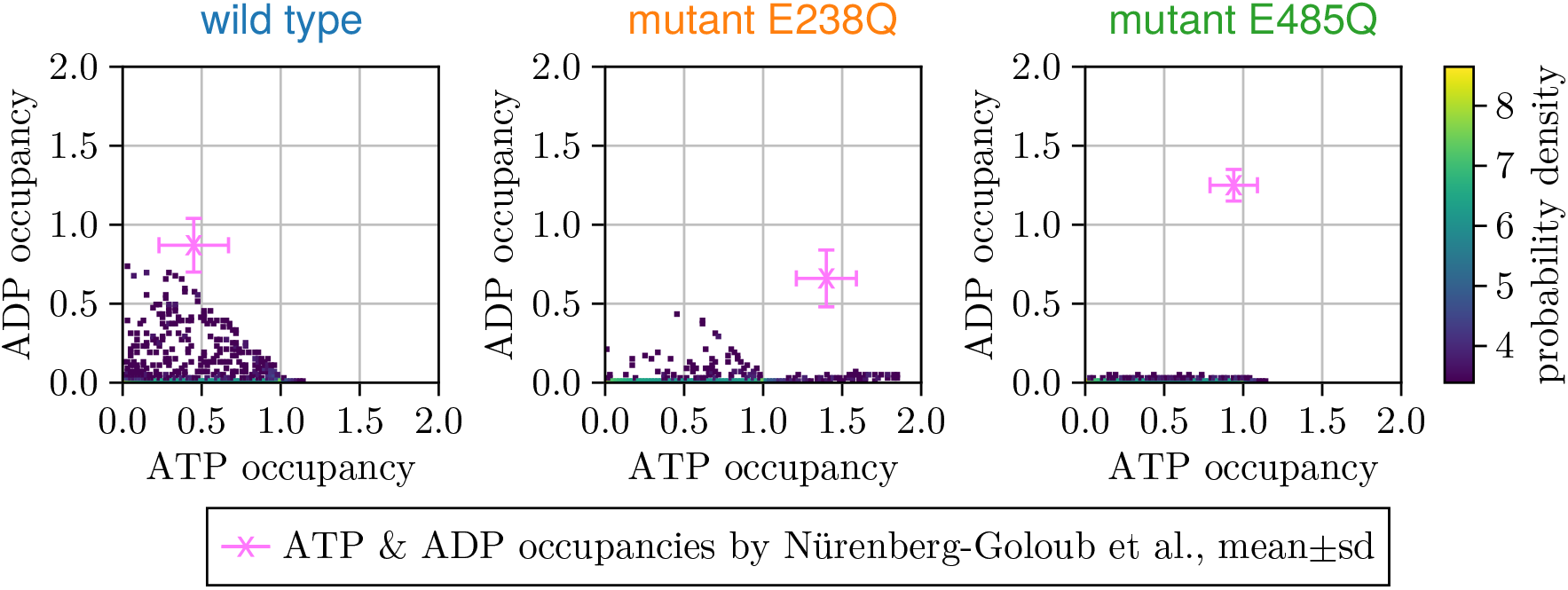
Ligand occupancies. Joint posterior probability distributions of single-turnover ATP and ADP occupancies are shown by color for wild type (left), mutant E238Q (middle) and mutant E485Q (right); the measured ADP and ATP occupancies^26^ (pink cross) and respective experimental uncertainties are shown for comparison.

As an alternative explanation we considered the fact that, due to the steady state charac ter of the experiments that were used to inform the Bayesian posterior, no kinetic information was available for the training of our Markov models. In contrast, the occupancies measured after 30 s do depend on the kinetics of the system and, hence, this disagreement is no sur prise. To test this idea, we included this kinetic information within the likelihood function and generated a new new Bayes sample, which, however, was still inconsistent with the observed non-equilibrium experiments (data not shown).

## 3 Conclusion

The ATPase ABCE1 has two nearly identical nucleotide-binding sites, both in terms of sequence and structure, suggesting that each nucleotide-binding site contributes equally to the overall ATP turnover rate. Recently, two mutants have been studied^7^to probe this near symmetry, each rendering one of the two nucleotide-binding sites ATP-hydrolysis defective. Only one of these mutants (E238Q) shows the expected twofold decrease of ATP turnover rate, whereas, quite unexpectedly, defectiveness of the other nucleotide-binding site (mutant E485Q) results in a staggering tenfold increase. A straightforward explanation would be an allosteric communication in terms of a direct interaction between the two sites, e.g., that the presence of ATP or ADP in one site strongly affects the binding affinity or hydrolysis rate of the other. Because for the ABCE1 system at hand this explanation is rather unlikely due to the high level of symmetry, we here asked whether the observed striking kinetic asymmetry between the two mutants can be explained also without any such direct allosteric interaction.

To answer this question, we described the combined conformational and chemical kinetics of ABCE1 by Markov models which, by construction, are thermodynamically consistent, but lack any direct communication between the two nucleotide-binding sites. As ‘minimal models’, our Markov models comprised 13 separate Markov states (each representing a combination of open and closed conformations with empty, ADP, and ATP occupation of each nucleotide-binding site) and physicochemically plausible transitions between these states. The respective transition rate coefficients were then determined via Bayesian inference using the measured ATP hydrolysis kinetics. In particular, a large Bayesian posterior sample of Markov models with specific transition rate coefficients for each transition was generated, yielding a posterior probability to agree with the measured data for each of these models. This posterior served to identify a subset of all possible Markov models that agrees with the observed ATP hydrolysis kinetics within experimental error, including the observed kinetic asymmetry. The fact that such a subset of Markov models exists demonstrates that, indeed, a direct allosteric communication between the nucleotide-binding sites is not required to explain the observed kinetic asymmetry.

This is the main result of this study. Because, obviously, an enzyme with two *fully independent* nucleotide-binding sites cannot exhibit the observed hyperactivity, we conclude that it is caused by the concerted opening and closing motion of the two protein domains. This conformational dynamics provides some communication between the two nucleotide-binding sites—albeit very indirectly—and was previously not considered a suitable explanation for the observed asymmetry or any allostery in general.

Closer inspection of the identified Markov models provided mechanistic insight into how the asymmetric ATP hydrolysis kinetics can be achieved. The most striking property, shared by almost one fifth of all Markov models that quantitatively agree with experiment and thus reproduce the observed asymmetry, is that in the wild type two separate reaction cycles contribute to the overall ATP turnover rate. Unexpectedly, both involve hydrolysis in NBSI, such that by far most of the ATP hydrolysis occurs within NBSI. While both cycles contribute about equally to the overall hydrolysis turnover rate, their population differs drastically by two to three orders of magnitude. In fact, the open ABCE1 conformation with ATP-loaded NBSI and ADP-loaded NBSII is by far the most populated Markov state. This population asymmetry is compensated by a corresponding rate coefficient asymmetry for the two cycles, resulting in similar fluxes, each contributing ca. 50 % to the overall hydrolysis rate. Crucially, ATP hydrolysis within NBSII connects these two cycles, thus creating a ‘drainage’ that creates this population asymmetry by steadily depleting the sparsely populated cycle in favor of the highly populated one. By blocking this drainage in the mutant E485Q, a fraction of the dominant population shifts towards the low-populated cycle, thus enhancing its contribution to the overall hydrolysis rate from 50 % to about 10-fold.

Using steady-state and new non-equilibrium measurements, we performed two rounds of cross validation aiming to check if our Markov model accurately describes the ATP hydrolysis kinetics of the ABCE1 wild type and the two mutants that knock out one of the active sites each. While the first cross validation showed consistency with the new data, the second did not. One possible reason is that our minimal 13-state Markov model is too small to capture all aspects of the complex ABCE1 ATP hydrolysis kinetics. Specifically, a half open conformation may be relevant, and it may be necessary to extend our Markov model accordingly to achieve quantitative agreement also with the non-equilibrium measurements. Of course, in addition to the mechanism postulated here, a direct effect of the occupancy of one nucleotide-binding site on the binding affinity or ATP hydrolysis rate of the other (e.g., of electrostatic nature), may exist and may be required to resolve the remaining discrepancies.

To narrow down which extension of our minimal model not only resolves the observed discrepancy, but also provides a description of the underlying mechanism, future experiments may help to discriminate between reaction types and to further restrict the parameter space. Recent advances with nanoaperture optical tweezers^52^, which enable time-resolved^53^ observation of conformational^54^ and binding dynamics^55^ at the single-molecule level, have promising potential to provide the required additional insight.

In summary, and independent of the above caveats regarding the proper modeling of ABCE1, this study revealed an unexpectedly complex behavior of Markov models, which can provide a thermodynamically consistent description of equilibrium, steady-state and non-equilibrium behavior of proteins that achieve their function through a tight interaction between conformational motions and chemical reactions. Even for systems as simple as the ATPase ABCE1, which comprised only two active sites and only two conformers, the resulting complexity of an 13-state minimal Markov model gives rise to — and can explain — quite counter-intuitive behavior. Specifically, using this type Markov model, our study demonstrates that the measured striking ATP hydrolysis kinetics asymmetry of ABCE1 does not necessarily require any direct allosteric interactions between the two nucleotide binding sites (nor does it rule it out, though); rather, this asymmetry can be explained solely in terms of concerted opening/closing conformational motions of this dimer. Notably, the 10-fold enhanced overall ATP hydrolysis rate upon *removal* of the ‘drainage’ represents a striking example of the so-called Braess’ paradox^56^, which in its original formulation states that removal of a road can enhance traffic throughput.

## 4 Methods

Markov models We use time-continuous Markov models to describe ABCE1 as a set of discrete states with memory-free transitions between them (Figure 2). The probability (per second) of such a transition from state *i* to state *j* is determined by its respective transition rate {*k*_*ji*_} ∈ ℝ_≥0_ The probability of ABCE1 to be in a state *i* at a given time *t* is given by *X*_*i*_(*t*), and, given an initial distribution ***x***(0), the time evolution of the probabilities ***x***(*t*) is determined by the master equation,

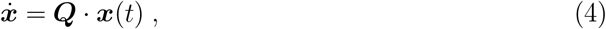

a system whose transition rate matrix ***Q*** contains the rate coefficients *k*_*ji*_ and diagonal elements 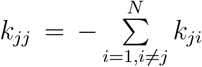, where *N* is the number of states. Note that *k*_*ji*_ are referred to as transition rates in the context of Markov models, but hereafter we refer to them as transition rate coefficients *k*_*ji*_ to be consistent with chemical terminology. For irreducible and aperiodic Markov models, the probabilities *x*(*t*) converge towards unique stationary or steady state probabilities ***π*** for *t* → ∞, such that

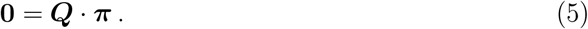

The state state probabilities ***π*** were calculated by solving the system of linear equations 5 without integration of the master equation 4^57^. Given ***x***(*t*), the net flux of a transition, i.e., how much probability (per second) is transferred between states *i* and *j*, is *k*_*ji*_*x*_*i*_ − *k*_*ij*_*x*_*j*_, where the signs are chosen such that the net flux is positive if ATP is hydrolyzed.

The linearity of the equations 4 and 5 implies that Markov models are inherently limited to describing first order reactions. In order to also include the higher-order reactions of ATP-, ADP-, and P_*i*_-binding within the Markovian framework, and following the steady state assumption^58^, we assume that the change of these concentrations is negligible at the timescale of the hydrolysis cycle and include these concentrations as constant factors of the respective transition rate coefficients in matrix ***Q***, for example, the transition rate coefficient for ATP-binding was chosen as *k*_*J*_ [ATP].

As discussed in the Results section, we further described ADP-binding and Pi-binding by a single rate-limiting transition with effective transition rate coefficients. To ensure that the free energy difference along each closed cycle including a net ATP hydrolysis still equals the Gibbs free energy of ATP hydrolysis 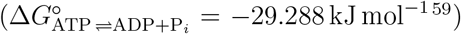, for ther modynamic consistency, the transition rate coefficient for ADP + P_*i*_-binding was chosen as *k*_*H*_[ADP][P_*i*_].

As also discussed in the Results section, we parametrized our Markov models by 17 free energy parameters, of which 7 were free energy differences (free energy difference associated with the conformational change Δ*G*_open→closed_, two free energy differences between ATP and ADP bound in the closed state Δ*G*_ATP→ADP, NBSI_ and Δ*G*_ATP→ADP, NBSII_, and four free binding energy differences Δ*G*_ATP binding, NBSI_, Δ*G*_ADP unbinding, NBSI_, Δ*G*_ATP binding, NBSII_, and Δ*G*_ADP unbinding, NBSII_) and 10 were forward free energy barriers (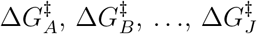 one for each transition A,B,….,J).

For each experiment the transition rate coefficients were scaled to the experimental temperature in accordance with transition state theory^46^.

### Description of steady state experiments

The measurements of limiting ATP turnover rate *k*_*cat*_, Michaelis-Menten constant *K*_*M*_, and ATP occupancy were assumed to be performed under steady state conditions^7^. Therefore, in order to calculate the posterior (equation 2, these observables had to be calculated from equation 5.

Based on the steady state solution *π* of equation 5, the ATP turnover rate was calculated as the sum of the net fluxes of all ADP-unbinding transitions for ATP concentrations between 10^−9^ and 10^2^ mol*/*L in logarithmic steps of 10.

The saturating ATP turnover rate *k*_vmax_ was calculated as the ATP turnover rate at a concentration of 10^2^ mol*/*L ATP.

The Michaelis-Menten constant was calculated from the ATP turnover rates determined by linear interpolation from two concentrations for which the turnover rates were close to *k*_*vmax*_*/*2. These two concentrations were determined from the two closest concentrations found for the above logarithmic grid by calculating ATP turnover rates for 18 additional logarithmically spaced concentrations between those, and again selecting those two closest to *k*_*vmax*_*/*2. This approximate calculation was chosen over more accurate ones because of the need for high computational efficiency for the extensive Bayes-sampling.

To construct the transition rate matrix *Q* of equation 5, ADP and P_*i*_ concentrations had to be chosen. The initial experimental ADP and P_*i*_ concentrations were zero^7^, however, due to ATP hydrolysis by ABCE1, these concentrations were on the order of the ABCE1 concentration during the experiment. To account for this increase, we selected the ADP and P_*i*_ concentrations to be 100-fold the median experimental ABCE1 concentration (5 *×* 10^−6^ mol L^−1^)^7^, assuming multiple completed ATP hydrolysis cycles of ABCE1. We opted for this more rudimentary estimate assuming that deviations from these ADP and P_*i*_ concentrations would exert a negligible influence on the calculated observables, which is consistent with the finding that ADP-/P_*i*_-binding is not identified as the rate-limiting step in any Markov model.

All ATP/ADP occupancies were calculated from the steady state population via 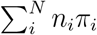, where *n*_*i*_ are the respective ATP/ADP occupation numbers *{*0, 1, 2*}*. To estimate the ligand concentrations at the time of the measurement, we used the integrated Michaelis-Menten kinetics, i.e., assuming a steady-state at each point of time, and used the mean of concentrations between 4 min and 5 min.

#### Description of non-steady state experiments

The ATP and ADP occupancy measure ments by Nürenberg-Goloub et al. were performed under single-turnover conditions with a 2:1 ratio of ATP to ABCE1^26^ and, therefore, the above steady-state assumption does not hold. We therefore resorted to numerical integration of equation 4, which in this case become third order, with initial values [ABCE1] = 0.3 *×* 10^−6^ mol*/*L, [ATP] = 0.6 *×* 10^−6^ mol*/*L, and [ADP] = [P_*i*_] = 0 mol*/*L. The numerical integration was performed using the Rodas4P^60^ algorithm with a maximal time step of 0.1 s, relative tolerance 10^−6^, and 10^6^ maximum steps, for a duration of 30 s, to obtain *x*(*t* = 30 s). The initial conditions assumed were that all ABCE1 is in an open conformation and both nucleotide-binding sites are empty.

#### Bayesian approach

For the calculation of the Bayesian posterior, we used a uniform prior for all 17 free energy parameters with boundaries chosen separately for chemical reactions, conformational transitions, and un-/binding events following physical plausibility (summarized in Table 1). To facilitate implementation, these boundaries were defined in terms of transition rate coefficients rather than free energies. For thermodynamic consistency, only those Markov models that satisfy detailed balance were considered.

**Table 1:**
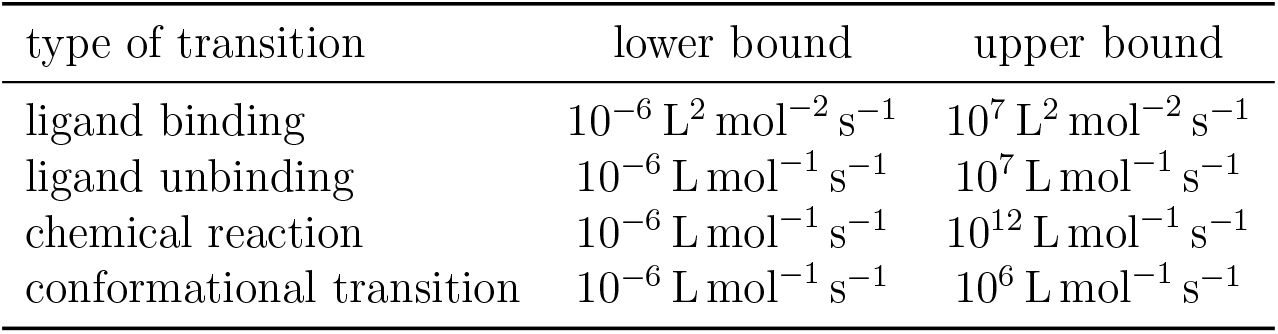
Summary of prior boundaries used for Bayes sampling. The upper limit for chemical reactions was assumed to be faster than that of binding/unbinding events or conformational transitions.

| type of transition | lower bound | upper bound |
| --- | --- | --- |
| ligand binding | $10^{-6} \text{ L}^2 \text{ mol}^{-2} \text{ s}^{-1}$ | $10^7 \text{ L}^2 \text{ mol}^{-2} \text{ s}^{-1}$ |
| ligand unbinding | $10^{-6} \text{ L mol}^{-1} \text{ s}^{-1}$ | $10^7 \text{ L mol}^{-1} \text{ s}^{-1}$ |
| chemical reaction | $10^{-6} \text{ L mol}^{-1} \text{ s}^{-1}$ | $10^{12} \text{ L mol}^{-1} \text{ s}^{-1}$ |
| conformational transition | $10^{-6} \text{ L mol}^{-1} \text{ s}^{-1}$ | $10^6 \text{ L mol}^{-1} \text{ s}^{-1}$ |

For the Bayes-sampling (equation 2), we used a replica-exchange Markov-chain Monte Carlo algorithm with a slice sampler for local exploration with 8 chains of 32 768 Markov models each using the Julia package Pigeons^47^. To provide a quantitative assessment of convergence supplementary Table S2 lists 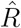-values^61^ of each chain and free energy parameter, in addition to the qualitative discussion of convergence in the Results section. 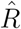-values close to 1 indicate sufficient convergence of most chains with slightly worse convergence of individual chains, for example, chain 2 of 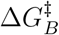 with 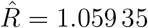

Rate-limiting step To calculate the rate-limiting step, we used backpropagation to calculate the derivative of the overall ATP turnover rate with respect to each unique transition rate (20) of a Markov model.

## 5 Acknowledgments

The authors thank Elina Nürenberg-Goloub, Yixin Chen, and Nicolai Kozlowski for helpful discussions. We thank Lars V. Bock for proofreading and providing valuable comments on the manuscript. H.G. and M.S. were supported by the Max Planck Society, M.S. by the International Max Planck Research School for Physics of Biological and Complex Systems, and R.T. by the German Research Foundation (DFG, TA157/15-1).

