## Supplementary Material for "Braess’ Paradox in Enzyme Kinetics: Asymmetry from Population Balance without Direct Cooperativity"

Table S1: Definition of transition rate coefficients in Figure 2. Listed are Markov transitions (left column), which describe either conformational transitions, binding reactions, or chemical reactions (right column). Occupations are indicated for NBSI/NBSII.

| transition rate coefficients | | Markov transitions | |
|---|---|---|---|
| forward | backward | forward | backward |
| $k_A$ | $k_{-A}$ | open → closed, ATP/ATP | closed → open, ATP/ATP |
| $k_B$ | $k_{-B}$ | ATP → ADP, NBSI | ADP → ATP, NBSI |
| $k_C$ | $k_{-C}$ | ATP → ADP, NBSII | ADP → ATP, NBSII |
| $k_D$ | $k_{-D}$ | closed → open, ADP/ATP | open → closed, ADP/ATP |
| $k_E$ | $k_{-E}$ | closed → open, ADP/ADP | open → closed, ADP/ADP |
| $k_F$ | $k_{-F}$ | closed → open, ADP/ADP | open → closed, ADP/ADP |
| $k_G$ | $k_{-G}$ | ADP unbinding, NBSI | ADP binding, NBSI |
| $k_H$ | $k_{-H}$ | ADP unbinding, NBSII | ADP binding, NBSII |
| $k_I$ | $k_{-I}$ | ATP binding, NBSI | ATP unbinding, NBSI |
| $k_J$ | $k_{-J}$ | ATP binding, NBSII | ATP unbinding, NBSII |

Table S2: Quantitative per chain Bayes sampling convergence assessment. Listed are $\hat{R}$-values[61] per chain and per free energy parameter.

| parameter | chain 1 | chain 2 | chain 3 | chain 4 | chain 5 | chain 6 | chain 7 | chain 8 |
|---|---|---|---|---|---|---|---|---|
| $\Delta G^{\ddagger}_{A}$ | 1.00747 | 1.00166 | 1.03971 | 1.00049 | 1.00634 | 1.00095 | 1.00589 | 1.00019 |
| $\Delta G^{\ddagger}_{B}$ | 1.00175 | 1.05935 | 1.00743 | 1.00177 | 1.00017 | 1.01509 | 1.01703 | 1.00976 |
| $\Delta G^{\ddagger}_{C}$ | 1.00135 | 1.00011 | 1.02143 | 1.0034 | 1.01301 | 1.01833 | 1.00405 | 1.01165 |
| $\Delta G^{\ddagger}_{D}$ | 1.00073 | 1.00082 | 1.00522 | 1.00279 | 1.00028 | 1.00254 | 1.00032 | 1.02118 |
| $\Delta G^{\ddagger}_{E}$ | 1.00539 | 1.00084 | 1.00183 | 1.0012 | 1.01355 | 1.01616 | 1.00363 | 1.00088 |
| $\Delta G^{\ddagger}_{F}$ | 1.00219 | 1.03133 | 1.00286 | 1.00016 | 1.00281 | 1.00313 | 1.0065 | 1.00103 |
| $\Delta G^{\ddagger}_{G}$ | 1.01779 | 1.00744 | 1.00355 | 1.00139 | 1.01159 | 1.04002 | 1.00034 | 1.03073 |
| $\Delta G^{\ddagger}_{H}$ | 1.00096 | 1.00381 | 1.00149 | 1.0012 | 1.00018 | 1.00131 | 1.00431 | 1.02721 |
| $\Delta G^{\ddagger}_{I}$ | 1.00005 | 1.00221 | 1.00047 | 1.00053 | 1.00025 | 1.00273 | 1.00004 | 1.00168 |
| $\Delta G^{\ddagger}_{J}$ | 1.00872 | 1.00485 | 1.00949 | 1.00038 | 1.00822 | 1.00798 | 1.00082 | 1.00445 |
| $\Delta G_{\text{ATP binding, NBSI}}$ | 1.01087 | 1.01359 | 1.00862 | 1.0001 | 1.01316 | 1.02756 | 1.01616 | 1.01208 |
| $\Delta G_{\text{ADP unbinding, NBSI}}$ | 1.00381 | 1.00191 | 1.00123 | 1.00028 | 1.00568 | 1.00789 | 1.00424 | 1.00298 |
| $\Delta G_{\text{ATP binding, NBSII}}$ | 1.00542 | 1.0072 | 1.01108 | 1.00041 | 1.00749 | 1.00218 | 1.00056 | 1.00089 |
| $\Delta G_{\text{ADP unbinding, NBSII}}$ | 1.00615 | 1.00132 | 1.01301 | 1.00063 | 1.00114 | 1.00431 | 1.00867 | 1.00044 |
| $\Delta G_{\text{open}\rightarrow\text{closed}}$ | 1.00065 | 1.01764 | 1.01318 | 1.00089 | 1.00436 | 1.00556 | 1.00448 | 1.00916 |
| $\Delta G_{\text{ATP}\rightarrow\text{ADP, NBSI}}$ | 1.00004 | 1.01081 | 1.00268 | 1.006 | 1.00248 | 1.00097 | 1.00358 | 1.02105 |
| $\Delta G_{\text{ATP}\rightarrow\text{ADP, NBSII}}$ | 1.00269 | 1.0014 | 1.01869 | 1.00165 | 1.00112 | 1.01113 | 1.00886 | 1.00224 |

Figure S1: Bayes sampling convergence assessment. Shown are sampled posterior probability densities of barrier heights $\Delta G^{\ddagger}$ and free energy differences $\Delta G$ for all transitions between Markov states. Histograms for 8 individual Markov Chains (thin black lines) as well as for all 8 chains combined (thick red lines) are shown. The inset (bottom right) lists $\hat{R}$-values for quantitative assessment of convergence.

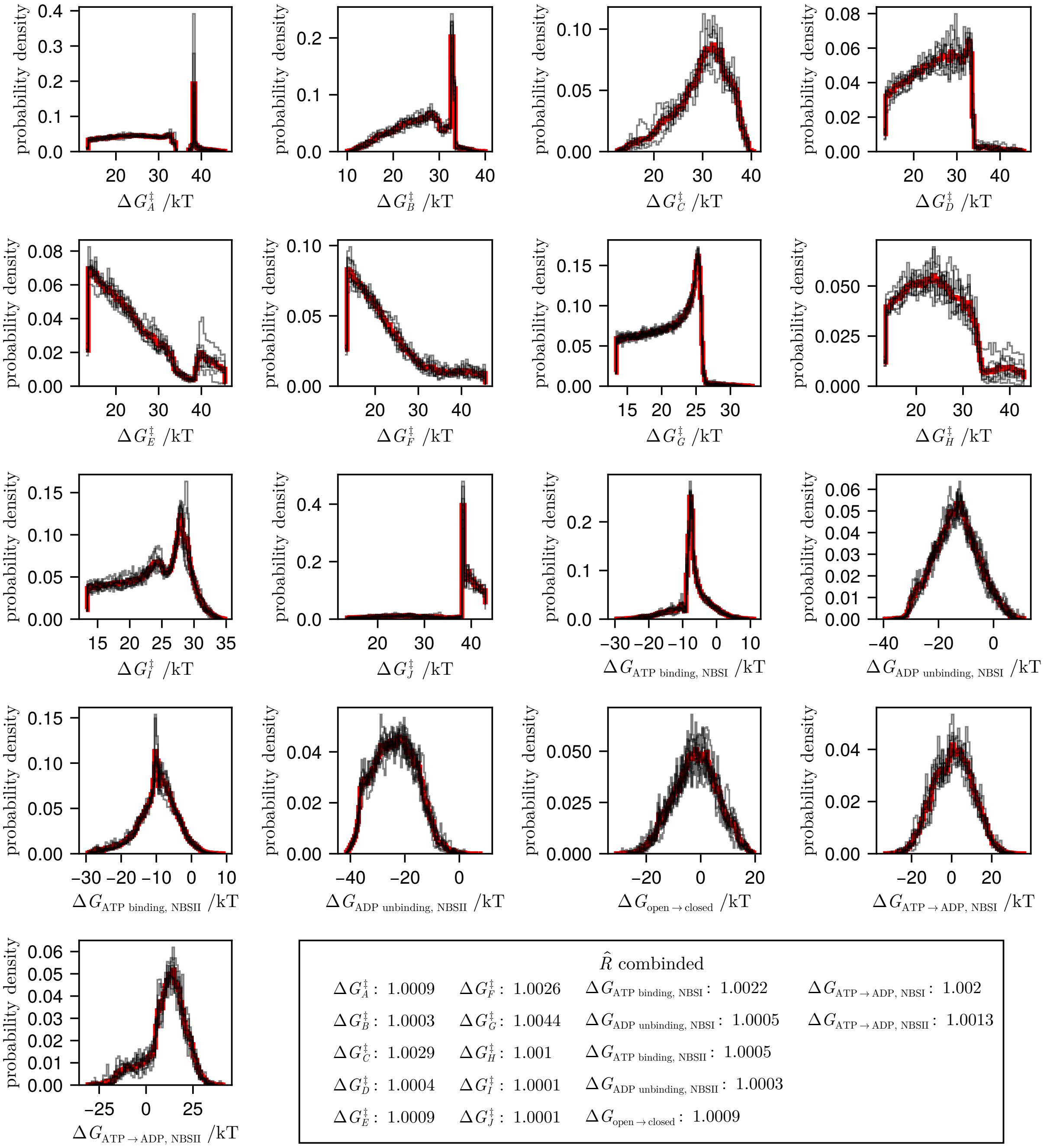

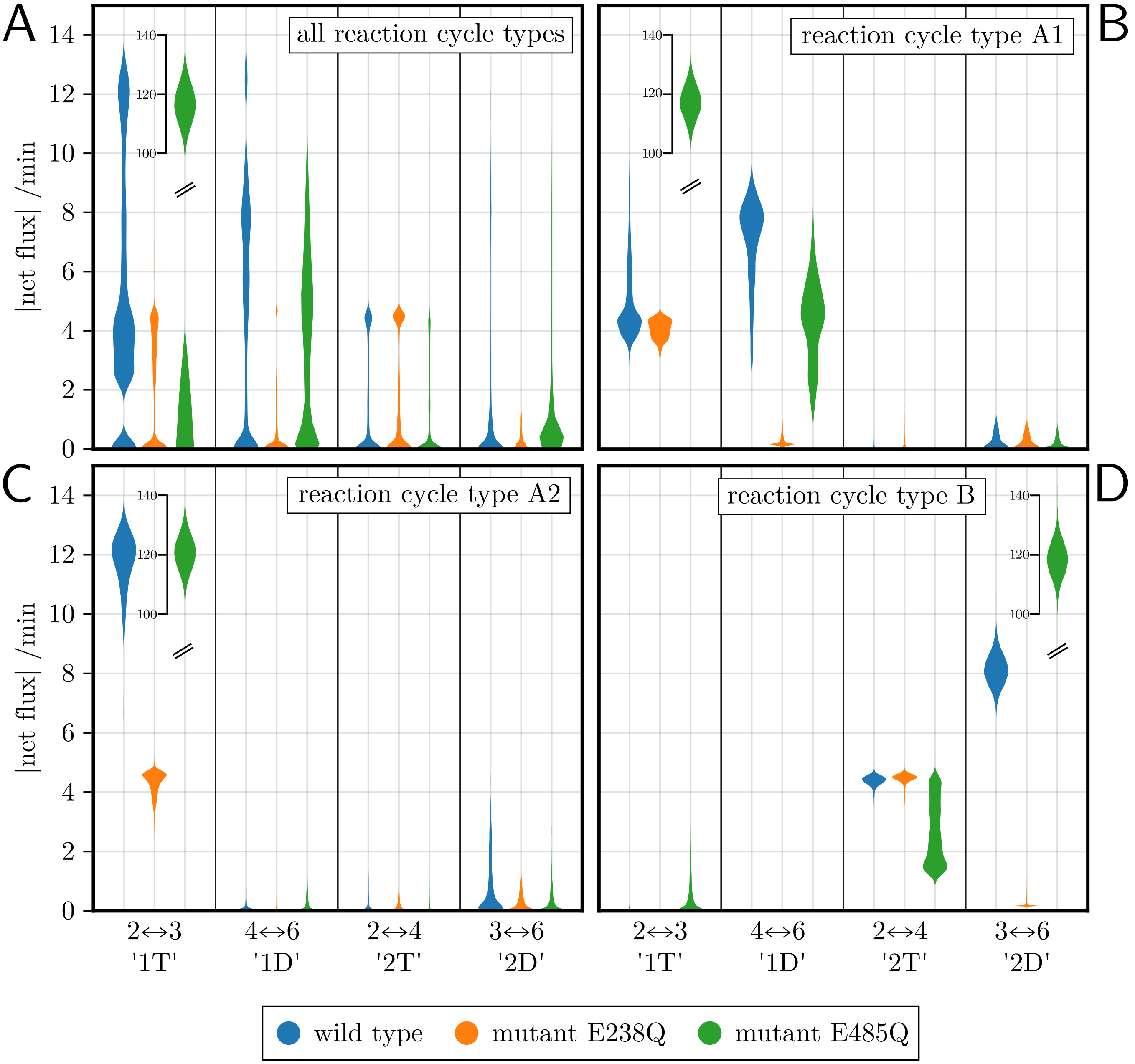


Figure S2: Net flux distributions, separated by ATP hydrolysis transitions. Shown are smoothed histograms of absolute net fluxes within Markov models for (A) all reaction cycles or (B-D) one of the three most probable reaction types (cf. Figure 4) for wild type and mutants (color). Annotations $i \leftrightarrow j$ indicate net fluxes between state $i$ and $j$, which are dominated by net fluxes through reaction cycles '1T', '1D', '2T', and '2D' (cf. Figure 4). Note the discontinuities in the axis to accommodate the high net fluxes observed in the E485Q mutant.

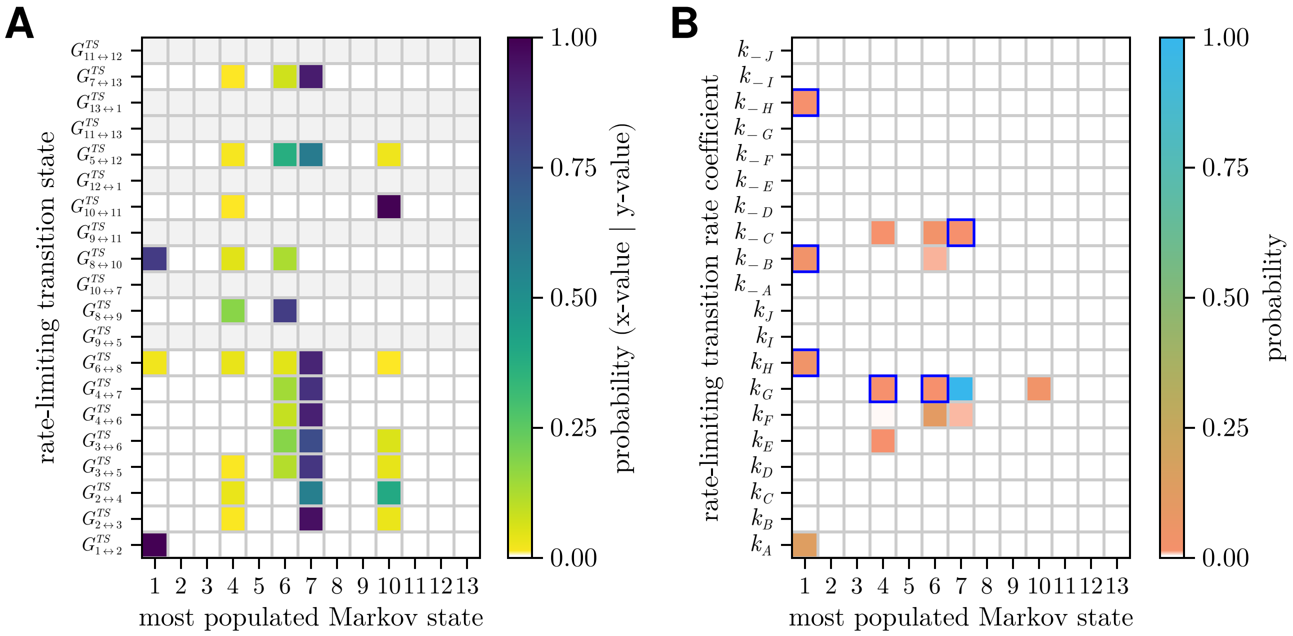


Figure S3: Comparison between most populated wild-type Markov states and kinetic determinants. (A) Shown are the conditional probabilities (color) of the most populated Markov state given each transition state to be rate-limiting. $G^{TS}_{i \leftrightarrow j}$ is the free energy of the transition state between state $i$ and $j$. For transition states or transition rate coefficients that were not rate-limiting in any Markov model, the rows are empty (gray). (B) Shown are the probabilities of each combination of rate-limiting transition rate coefficient and most populated Markov state. Blue rectangles indicate combinations where the rate-limiting transition state does not belong to an outgoing transition of the most populated Markov state.
